# Combining structural modeling and deep learning to calculate the *E. coli* protein interactome and functional networks

**DOI:** 10.1101/2025.05.07.652715

**Authors:** H. Zhao, C. Velez, A. Navarene, A. Saha, J. Feldman, J. Skolnick, D. Murray, B. Honig

## Abstract

We report on the integration of three methods that are computationally efficient enough to predict, on a proteome-wide scale, whether two proteins are likely to form a binary complex. The methods include PrePPI, which uses three-dimensional structure information as a basis for predictions, Topsy-Turvy which analyzes sequences using a protein language model, and ZEPPI which uses evolutionary information to evaluate protein-protein interfaces. We demonstrate how these methods can be integrated and validate the performance of the integrated method and its separate components at predicting *E. coli* protein-protein interactions through testing on the HINT high-quality literature-curated database of binary interactions. The integrated method identifies more high-confidence (FPR ≤ 0.001) interactions (∼20K) than any of the component methods. The AF3Complex algorithm was used to predict the structures of 400 protein-protein interactions, and 78% of the integrated method predictions resulted in models deemed accurate by the AF3Complex evaluation score. Notably, essentially all AF3Complex models have at least partially overlapping interfaces with PrePPI models of the complexes. Finally, we clustered the high-confidence *E. coli* interactome and obtained 385 subnetworks which have high functional coherence defined by enrichment of Gene Ontology Biological Process terms, thus, illustrating that our methods which contain no explicit functional information provide biologically meaningful protein interactions. Biological insights derived from the subnetworks, including the annotation of proteins of unknown function, are discussed in detail. Overall, independent validations support the accuracy of the comprehensive *E. coli* interactome we have presented.

## Introduction

Predicting whether two proteins interact and, where relevant, building a model of the complex they form have become problems of great interest. This is due both to the multitude of biological applications that the knowledge of protein-protein interactions (PPIs) enables and to major developments in computational approaches, particularly those based on AlphaFold (AF)^1^. The AlphaFold group at Google DeepMind has made models available for over 200 million proteins from multiple organisms^1^ (the AF2 database – AFDB) and, in addition, has developed methods to predict the structures of two or more interacting proteins^2^. The wide adoption of AF-based methods by the biological community has been remarkable with models of individual proteins downloaded from AFDB and of complexes, based on user-friendly programs such as AlphaFold Multimer^3^ and AF3^2^, becoming common features in the biological literature.

Our interest in this work is the prediction of PPIs on a proteome-wide scale, involving for example, ∼200 million PPIs for human, ∼18 million for yeast and ∼9 million for *E. coli*. The computational cost of carrying out AF-based calculations for so many complexes is prohibitive, especially when accounting for multi-domain proteins where treating each domain as a separate entity will increase these numbers by over an order of magnitude. Nevertheless, the challenge of deriving proteome-wide interactomes is of great importance as they can reveal the details of complex PPI networks that control cellular function. High throughput experimental methods, in particular, have made major advances in large scale PPI detection but have not yet reached the scale of complete interactomes and, further, do not reveal structural information.

Computational approaches have also made significant progress on the proteome-wide prediction of PPIs and a wide range of strategies have been deployed^4^. Baker and colleagues have developed a strategy involving filtering full interactomes using co-evolution-based methods which significantly reduces the number of PPIs to be considered and then applying structure-informed methods to evaluate the PPIs that remain. In their original paper on the *E. coli* proteome, protein-protein docking was used to evaluate the ∼22,000 PPIs that passed the co-evolution filter and, finally, 804 high confidence PPIs were identified for further analysis^5^. In more recent, post-AF studies on yeast^6^ and humans^7^, putative PPIs were first filtered based solely on co-evolution and further filtered with Rosetta fold after which AF was used to identify high confidence interactions. For humans, about 7,000 high confidence PPIs (expected precision >90%) were identified with the *de-novo* pipeline. The end point of these studies has been the identification of extremely high confidence interactions, some of which were not reported in experimental databases. However, they have not been implemented to provide a proteome-wide PPI interactome that can then be used to describe functional subnetworks. PrePPI (Predicting Protein-Protein Interactions), which was developed with this goal in mind, adopts a very different strategy^8-10^.

The PrePPI pipeline begins with structural models of the isolated query proteins and searches for pairs of structurally similar proteins that form a complex in the Protein Data Bank (PDB)^11^. When a PDB complex is found, the query protein models are superimposed on the respective template chains, thus producing a homology model of the query complex. This model is then evaluated with an extremely efficient scoring function which considers only the quality of the structural alignment and the residues in the query interface that align with those in the template interface. PrePPI’s efficiency is such that it can score billions of models between interacting proteins and their individual domains. PrePPI uses Bayesian statistics to train and score models and, thus, properly chosen training and test sets are of crucial importance. The full version of PrePPI^9^ includes various sources of non-structural evidence, but here we consider only structural modeling, designated below as SM. Of note, PrePPI has been subject to a number of direct experimental tests beyond comparisons to various databases. Direct tests have included a series of pulldown experiments^9, 12^ and, most recently, the ability to predict proteins that are synthetically lethal to a constitutively activated mutant form of KRAS in experiments on mouse PDX models^13^.

It is important to emphasize that the quality of PrePPI models depends on the structural similarity of the query proteins to the PDB template proteins and that this similarity is reflected in the scoring function and, concomitantly, in the likelihood ratio (LR) associated with a particular PPI. Close homologs will yield good models while more distant homologs may yield complexes that produce only a weak signal (low LR) but may still be strong enough to provide meaningful evidence for the existence of a PPI. With the goal of further increasing the reliability of PrePPI predictions, we sought a way to integrate the PrePPI pipeline with independent sequence-based methods based on deep learning. This requires that we use a method that, like PrePPI, is fast enough, to predict PPIs on a proteome-wide scale. D-SCRIPT^14^ and Topsy-Turvy (TT)^15^, developed in the Berger and Cowen labs appear ideally suited for this purpose.

For a given PPI, we accomplish this integration with a Bayesian approach by multiplying LRs obtained from each source for a given PPI. These sources include: 1) PrePPI structural modeling scores (SM^LR^); 2) ZEPPI Z-scores derived from evolutionary analysis of PPI interfaces with respect to randomly generated interfaces^16^; and 3) TT interaction probabilities, TT^prob^, obtained from a protein language model and network topology considerations^15^. The ZEPPI Z-scores and TT probabilities are transformed into LRs, ZEPPI^LR^ and TT^LR^, through additional training and combined with SM^LR^ into an integrated score, Int^LR^.

We report and compare the performance of three approaches (SM^LR^, TT^LR^, and Int^LR^) obtained from training on human PPIs and applied to *E. coli K12*. We also evaluate the performance of TT^prob^ which was trained on PPIs annotated as physical in the STRING database^17^. We find that PrePPI and TT-based methods perform comparably, and, at high confidence, predict largely different sets of PPIs. The main effect of combining all evidence sources is to identify many more high confidence PPIs.

TT does not produce atomistic models for complexes while PrePPI produces models of varying quality largely dependent on the SM^LR^ associated with the prediction. Both methods can be used to identify PPIs that can potentially be well-modelled with AF-based methods. We have chosen 400 predicted PPIs spread over the three prediction methods (PrePPI, TT^prob^ and Int) and used the AF3Complex algorithm to construct models for each^18^. Built from AlphaFold 3, AF3Complex achieves superior performance on protein complex prediction thanks to its modified MSA generation algorithm, novel confidence metric, and self-selection of ligand and ion data. Moreover, thanks to the modified MSA generation algorithm in AF3Complex, the computationally expensive step of pairing MSAs is avoided, potentially saving a great deal of time and computational expenditure, making it well-suited for large-scale PPI prediction. We find that most predicted PPIs yield high confidence AF3Complex structural models, thus opening the door to a strategy involving fast proteome-wide evaluation of protein pairs followed by slower and more rigorous AF3Complex model building of the subset of those pairs most likely to form a binary complex.

Finally, we perform Markov clustering ^19^ to identify PPI subnetworks for each of the interactomes. Remarkably, *de novo* predictions of PPIs based on Int, TT, and PrePPI produce multiple sets of functionally coherent clusters. We provide an overview of a number of these clusters from the Int interactome and present AF3Complex models for some of the novel PPIs that are predicted to be cluster members.

## Results

### Performance comparison of different methods

Table 1 reports area under the Receiver Operating Character (ROC) and Precision Recall (PR) curves derived from SM^LR^, TT^prob^, TT^LR^ and Int^LR^ as evaluated by the 2021 HINT literature-curated high-quality binary database for *E. coli (*denoted here simply as HINT)^20^. TT^prob^ yields values based on the published TT algorithm^15^, which was trained on interactions in STRING defined as physical (denoted here as STRING-physical)^17^, while TT^LR^ is the result of additional training, also on the STRING database (see Methods). TT^LR^ performs slightly better than the TT^prob^ when tested on HINT demonstrating that the additional round of training has not diminished the performance of the published TT algorithm. The results in Table 1 also demonstrate that SM^LR^ exhibits somewhat improved performance relative to the two TT-based metrics, but it should be recalled that PrePPI was trained on the HINT database of literature-curated high-quality binary PPIs for human while both TT methods were trained on STRING-physical which includes non-direct interactions between proteins in the same multi-protein complex. Finally, Int^LR^ produces the best results suggesting the value of integrating different methods.

**Table 1.**
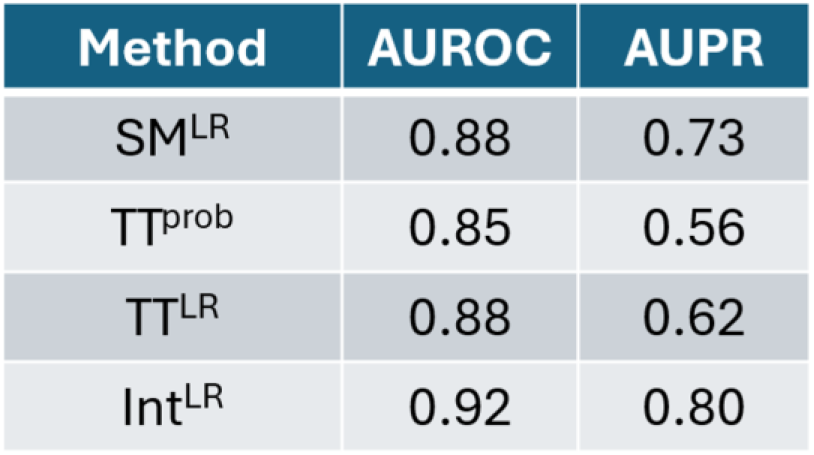
Evaluation metrics of PPI prediction methods. The areas under the Receiver Operating Characteristic and Precision Recall curves (AUROC and AUPR) were calculated using a 1:10 ratio of TP/TN.

Figure 1 displays ROC and PR curves corresponding to most of the values in Table 1. The TT curve is based on TT^prob^ to properly reflect the reported method. As can be seen from the ROC curves in the figure, SM^LR^ performs the best at very low FPR. At low FPR (Figure 1A) and low recovery rates (Figure 1B), SM^LR^ significantly outperforms TT^prob^ while Int^LR^ performs comparably to SM^LR^. The full SM^LR^ and TT^prob^ curves (Figure 1A, inset) are very similar as also reflected in the AUROC and AUPR values shown in Table 1. However, the AUROC and AUPR values are highest for Int^LR^ indicating again the value of combining the two methods. However, ROC and PR curves only tell part of the story.

**Figure 1.**
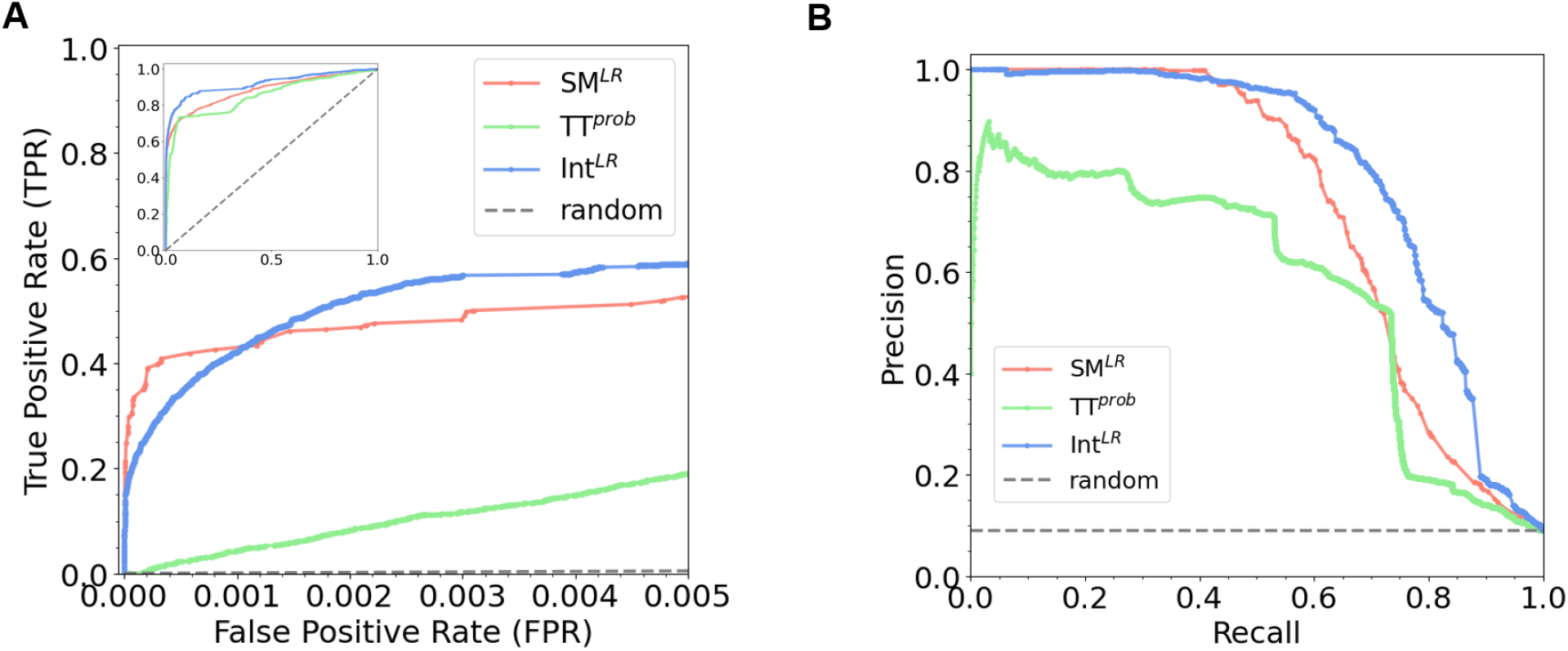
Evaluation of PPI prediction methods. A. Receiver Operating Characteristic (ROC) curve and B. Precision Recall (PR) curve as tested on the HINT high-quality literature-curated binary *E. coli* PPI set for Int (blue), SM^LR^ (red) and TT^Prob^ (green).

Figure 2 contains a Venn diagram for the number of PPIs predicted by SM^LR^, TT^LL^ and Int^LR^ at an FPR cutoff of ≤ 0.005. Of note, at this FPR cutoff TT^LL^ makes more predictions than SM^LR^ and there is relatively low overlap between the two sets of predictions indicating their complementarity. However, their combination results in an increase in coverage relative to either method alone and, notably, Int^LR^ provides 23,058 predictions found neither in SM^LR^ nor TT^LR^ alone (FPR≤0.005) illustrating the synergism between the different approaches. Overall, the main effect of method integration is a large increase in the number of high-confidence predictions.

**Figure 2.**
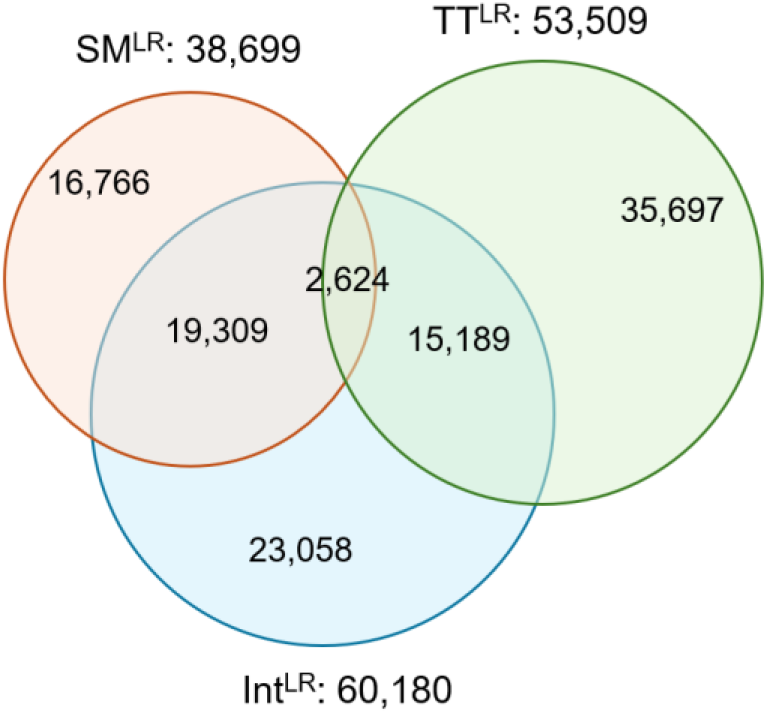
Overlap of PPIs from prediction methods. Venn Diagram for Int^LR^ (blue), SM^LR^ (red), and TT^Prob^ (green) tested on the HINT high-quality literature-curated binary *E. coli* PPI set. The numbers refer to PPIs associated with an FPR ≤ 0.005 for each ROC curve.

### AF3Complex models of predicted PPIs

AF3Complex^18^ was used to model 400 PPIs chosen from the PPIs predicted by Int with an FPR ≤ 0.001. The subset of 400 PPIs was filtered as described in Methods so as to focus on challenging cases (e.g. < 40% sequence identity between the query sequences and the respective chains in the PDB complexes chosen by PrePPI as modeling templates) and to represent a range of contributions of SM, TT and ZEPPI to the integrated score, Int^LR^. AF3Complex uses a predicted interface score, pIS, which focuses on the interfacial residues of a model, thus, improving structural modeling of predicted protein complexes. The suggested threshold for accurate AF3Complex models is pIS ≥ 0.38^18^ for 80% confidence predictions. As depicted in Supplementary Table S1A, AF3Complex produced scored models for 382/400 (96%) of Int PPIs, and, of these, 312/382 (82%) of the scored models meet the ≥ 0.38^18^ criterion for accurate models. Overall, 312/400 (78%) Int^LR^ predictions are associated with accurate AF3Complex models. For SM^LR^ (Figure 1, red lines), 284/400 (72%) predictions with SM^LR^ FPR ≤ 0.001 have accurate AF3Complex models. For TT^prob^ (Figure 1, green lines), 101/400 (25%) predictions with TT^prob^ FPR ≤ 0.001 have accurate AF3Complex models. However, TT uses a probability cutoff of 0.5 to determine if two proteins are predicted to interact^15^ and, based on this criterion, 291/400 (73%) predictions with TT^prob^ ≥ 0.5 have accurate AF3Complex models. That the integrated (Int) PPI prediction method yields the highest percentage (78%) of accurate models as determined by AF3Complex is a strong indication that many of our PPI binary predictions have identified true physical interactions.

Figure 3 provides a gallery of 12 high-scoring AF3Complex models (pIS > 0.5) for predictions from the FPR ≤ 0.001 region of the Int^LR^ ROC curve including a range of SM^LR^ and TT^prob^ values (Supplementary Table S1B). Supplementary Figure S1 contains the models in Figure 3 structurally superimposed with the respective PrePPI SM models. In half of the cases, the RMSD between the AF3Complex and PrePPI models is < 4Å, and in most (67%), the RMSD is < 6Å. However, in all cases the interfaces are reasonably well-aligned with interfacial residues being conserved on both sides of each interface (the last two columns of Supplementary Table S1B). In Figure S1 panels F, I, J, and K, the large RMSD values are driven mainly by changes in domain orientation. Overall, these results highlight the similarity between high confidence PrePPI and AF3Complex models. It is unclear at this point which models are more accurate, but this is likely dependent in part on the availability for good templates for PrePPI which does not optimize monomer structures as is often done so effectively by AF3.

**Figure 3.**
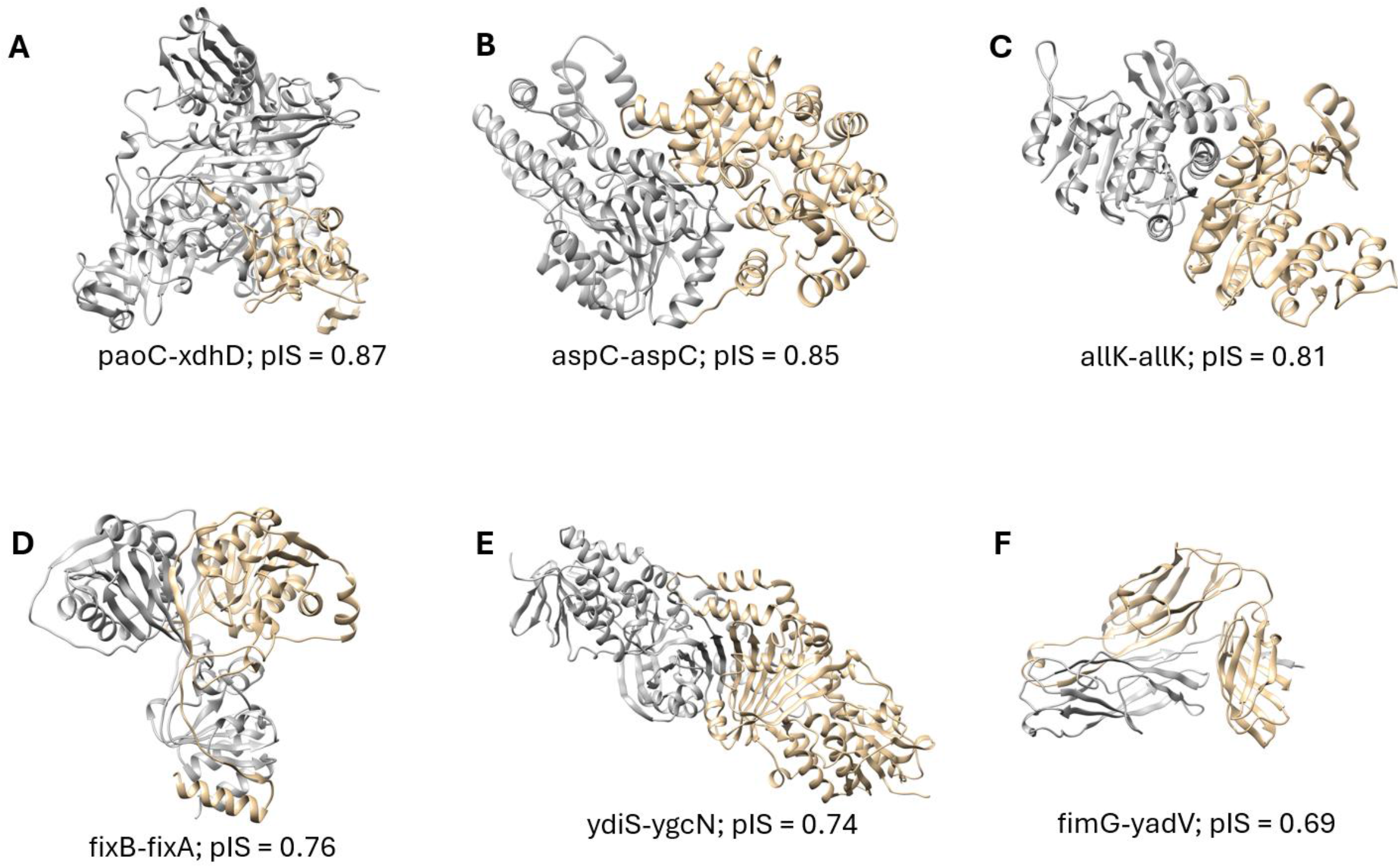
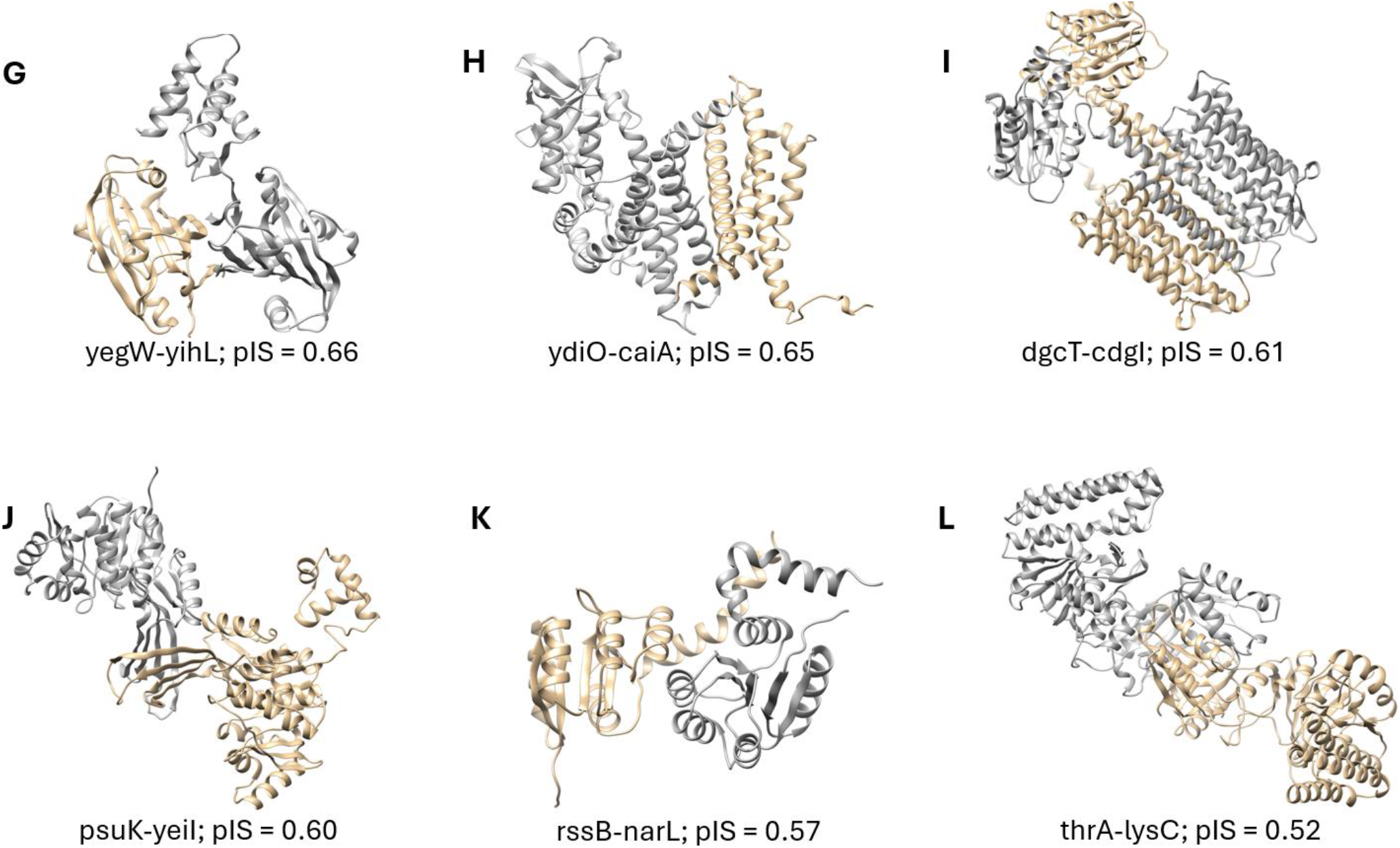
AF3Complex models for high-confidence Int^LR^ *E. coli* PPI predictions. Panels A through L depict models in backbone ribbon representation with one chain colored gray and the other colored gold. The PPIs were chosen from the FPR ≤ 0.001 region of the Int^LR^ ROC curve in Figure 1A (blue). pIS is the predicted Interface Score from AF3Complex. For simplicity, proteins are denoted by their gene names, and their UniProt IDs are provided in Supplementary Table S1B along with other details related to the predictions. Several models (panels A, F, H, and K) are discussed in the text. **A-F.** AF3Complex models for high-confidence Int^LR^ *E. coli* PPI predictions.

The AF3Complex models represent novel PPIs and known PPIs which have evidence from curated databases. For example, panel A displays a complex for the aldehyde oxidoreductase molybdenum-binding subunit protein PaoC and the hypoxanthine oxidase XdhD as mediated by the iron-sulfur subunit of XdhD (gold). PaoC is involved in purine nucleoside biosynthesis, whereas XdhD is involved in cellular detoxification. Both contribute to the synthesis of molybdenum cofactors^21^ which are essential for both pathways. Panel H is the AF3Complex model for the novel PPI between the acyl-CoA dehydrogenase, YdiO, and the crotonobetainyl-CoA reductase CaiA, which both function in fatty acid oxidation. The models in panels F and K are discussed in the context of functional clusters in the next section.

### Clustered PPIs yield functional subnetworks

At a False Positive Rate (FRP) ≤ 0.001 for the Int^LR^ network (Figure 1A), there are 20,523 protein-protein interactions (PPIs) among 3,219 proteins (73% of the *E. coli K12* proteome). The Int^LR^ network is comprised of 18,354 heterodimeric “edges” among 2,913 proteins (nodes). The network was visualized and analyzed in Cytoscape^22^. Network statistics are provided in Supplemental Table S2.A. Markov clustering as implemented in Cytoscape^19^ (with default parameters and edge weights set to 1 for all PPIs below an FPR threshold of 0.001) produced 558 clusters with 2 or more members (Supplemental Table S2.B). 139 proteins appear in no cluster, and 124/558 clusters (22%) contain 5 or more proteins with the largest cluster comprised of 1,632 PPIs among 124 proteins (Supplemental Table S2.C).

The clusters derived from the FRP ≤ 0.001 Int^LR^ interactome have high functional coherence (Figure 4). It is important to emphasize that these functional clusters are derived entirely based on our predicted Int^LR^ PPI interactome that contains no explicit functional information. The spectrum of biological processes assigned to the clusters is wide, encompassing terms including cell adhesion, regulation of DNA-templated transcription, phosphorelay signaling, anaerobic respiration, defense mechanisms, and a wide variety of transport processes and cellular component syntheses. Enriched Gene Ontology (GO)^23^ for individual clusters were calculated with clusterProfiler^24^, and terms with an adjusted p-value ≤ 0.05 were retained (Supplemental Table S2.D). 11,977 edges of the original 18,354 (65%) are preserved in the clustered network. 485/558 clusters (87%) are represented by at least one type of annotation, and 385 clusters (69%) are annotated with GO biological process (GO:BP) terms (Supplemental Table S2.E). The 385 GO:BP clusters contain 257 proteins that are not assigned GO:BP terms in the UniProt Knowledgebase (UniProtKB)^25^ (Supplemental Table S2.B). The occurrence of an unannotated protein in an annotated cluster suggests that the protein may participate in the same pathways and cellular processes in which the cluster is enriched. While the SM^LR^ and TT^prob^ interactomes (FPR ≤ 0.001) can be similarly clustered, the SM^LR^ clusters provide denser subnetworks and more statistically significant functional annotations (akin to Int^LR^ clustering) than the TT^prob^ clusters.

**Figure 4.**
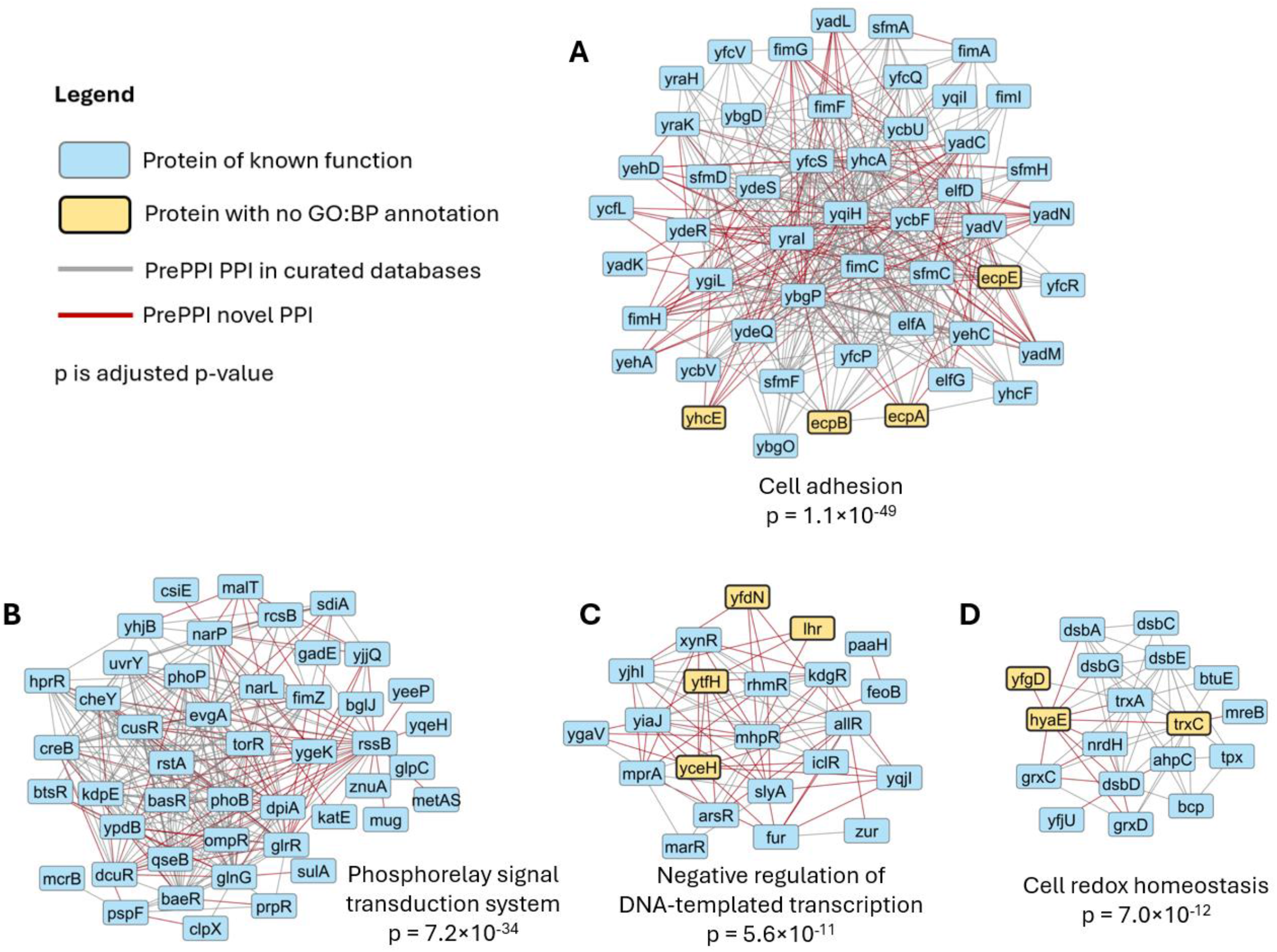
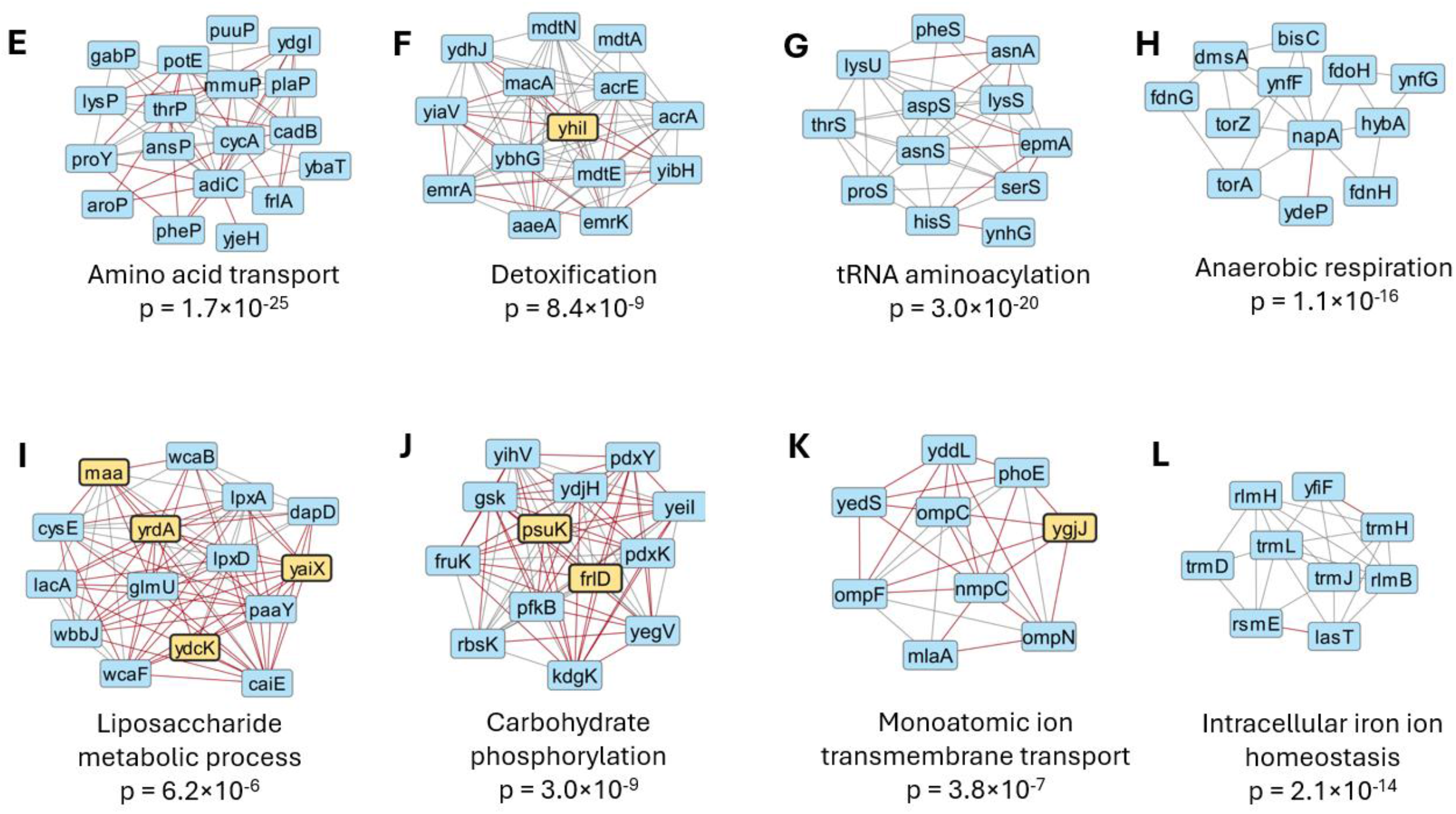
Subnetworks of the Integrated (Int) *E. coli* interactome. The PPI network comprised of Int^LR^ predictions (FPR ≤ 0.001) was clustered^19, 22^ and annotated with Gene Ontology (GO) terms^24^ as described in Methods. Panels A through L depict subnetworks comprised of a given cluster’s proteins (nodes) and interactions (edges) from Int^LR^. Blue nodes depict proteins associated with GO biological process (GO:BP) terms, and orange nodes depict proteins with no associated GO:BP annotation from UniProtKB. PPIs in curated databases are denoted by gray edges and novel PPIs are denoted by red edges. Below each cluster is its most statistically significant GO:BP term and that annotation’s adjusted p-value (p) (Supplementary Table S2.E). Clusters A through D are discussed in detail in the text, and Supplementary Table S2 provides complete clustering and annotation analysis results for the Int^LR^ interactome. **A-D**. Subnetworks of the Integrated (Int) *E. coli* interactome. **E-L**. Subnetworks of the Integrated (Int) *E. coli* interactome.

Supplemental Table S2.E contains top GO:BP terms for the 117 annotated clusters from the 124 clusters of five or more members. Examples of annotated clusters, some with proteins lacking GO:BP terms, are provided in Figure 4. For simplicity, proteins are denoted by their gene names and identified as necessary below. In the following, we elaborate on four clusters (Figure 4A-4D) which contain many novel PPIs. We also highlight potential annotations for proteins with no GO:BP annotation and provide AF3Complex models for novel PPIs to demonstrate how our clusters provide a source of functional annotation and hypothesis generation.

Figure 4A depicts a subnetwork highly enriched in proteins involved in “cell adhesion” with 33 of the 50 proteins having this GO:BP term, suggesting that the remaining 17 may also participate. Indeed, 11 of the 17 have the annotation “cell wall organization,” a related biological process that supports cell surface interactions. Of the four unannotated proteins, yhcE (the UPF0056 membrane protein YhcE) is a plasma membrane associated transmembrane protein predicted to interact with proteins involved in the GO:BP “cell wall organization” (fimC, yfcS, ybqP, yraI, and yqiH). The last steps of cell wall synthesis occur along the plasma membrane^26^, which suggests yhcE may participate in adhesion. The other three, ecpA (Common pilus major fimbrillin subunit EcpA), ecpB (Probable fimbrial chaperone EcpB), and ecpE (Probable fimbrial chaperone EcpE), are part of the operon that encodes the “pilus”, the hair-like cell-surface appendage that mediates cell adhesion^27^. Figure 3F depicts the AF3Complex model (pIS = 0.69) for the novel PPI between the fimbrial adhesion protein fimG (Protein FimG) and ydaV (Uncharacterized protein YdaV0), a chaperone for fimbriae assembly at the pilus^28^. ydaV facilitates the folding of fimG and the model provides a structural mechanism for this process. The analysis of this cluster provides not only novel annotations but also high-resolution structural models for novel interactions among cell adhesion proteins.

In Figure 4B, 72% of the PPIs in the subnetwork for “phosphorelay signal transduction system” are in databases of curated PPIs providing additional evidence for the functional coherence of our interactomes. Phosphorelays are two-component “intracellular signal transduction” systems that respond to environmental changes through a series of phosphorylation events involving histidine kinases and response regulators. One-third of the proteins in this cluster are part of a “protein-DNA complex” (GO:CC), including rssB (Regulator of RpoS), which directs the expression of genes involved in survival under stressful conditions, and narL (Nitrate/nitrite response regulator protein NarL), which regulates the expression of genes involved in anaerobic respiration. Figure 3K depicts the AlphaF3Complex model (pIS = 0.57) for the novel interaction between rssB (gray) and narL (gold) mediated by their receiver domains suggesting a means of interconnection of the two functions^29^.

The subnetwork in Figure 4C is involved in “negative regulation of DNA-templated transcription”. The four unannotated proteins – yfdN (Uncharacterized protein YfdN), lhr (Probable ATP-dependent helicase Lhr), ytfH (Uncharacterized HTH-type transcriptional regulator YtfH), and yceH (UPF0502 protein YceH) – interact with proteins associated with this GO:BP term. In addition, ytfH, which contains a transcriptional regulator domain, is predicted to make novel interactions with two essential proteins in this cluster: the transcriptional activator mhpR (DNA-binding transcriptional activator MhpR) and the transcriptional repressor allR (HTH-type transcriptional repressor AllR), which share a negative genetic interaction^30^.

The subnetwork in Figure 4D is based on the cluster annotated with “cell redox homeostasis.” trxC (Thioredoxin 2) is predicted to interact with trxA (Thioredoxin 1), and both have been shown to be part of the *E. coli* thioredoxin (Trx) system, which maintains thiol-based redox homeostasis^31^. trxC is also predicted to interact with proteins annotated as being localized to the “periplasmic space” (GO:CC), the oxidizing environment for cell redox homeostasis. Essentially all PPIs involving trxC in this subnetwork appear in curated databases (gray edges). Although trxC is annotated as a “cytoplasmic/cytosolic protein” (GO:CC), its interaction with periplasmic proteins indicates that it plays a role in cell redox homeostasis as well. Both hyaE (Hydrogenase-1 operon protein HyaE) and yfgD (Uncharacterized protein YfgD) are annotated with oxidoreductase activity (GO:MF). Novel PPIs between hyaE and periplasmic proteins involved in cell redox homeostasis suggest that it also plays a role in this process.

## Discussion

The number of possible pairwise combinations of proteins in the human proteome, if multiple isoforms are not considered, is approximately 200 million and on the order of billions when proteins are parsed at the domain level. Even high-throughput experimental methods^32-34^ cannot cover this range of possible PPIs so that computational approaches offer the most tractable approach to identify PPIs on a proteome-wide scale. AF-derived methods can produce accurate structural models of multimeric PPIs, but they are too computationally intensive for proteome-wide applications. PrePPI is computationally efficient enough for this purpose and has been shown to have an accuracy at least comparable to that of high throughput experiments^9, 10^, with additional confirmation through direct experimental tests^10, 12, 13^. Recently, protein language models have been used to derive sequence-based PPI prediction algorithms such as DSCRIPT and TT and, like PrePPI, are fast enough to be applied proteome wide^14, 15^. Moreover, since they do not rely on protein 3D structure, they can predict PPIs that are not accessible to PrePPI analysis. On this basis, we reasoned that the integration of PrePPI and TT would both improve prediction accuracy and increase the coverage obtained from PrePPI alone.

We have also added a new scoring function, ZEPPI, to evaluate PrePPI-predicted models^16^. Whereas PrePPI’s model evaluation is based on the comparison of predicted interfacial residues to those of a template, ZEPPI adds evolutionary analysis of contacting residues in the model itself and can improve the accuracy of PrePPI predictions. PrePPI uses a naïve Bayesian approach to combine evidence sources while ZEPPI provides Z-values and TT provides interaction probabilities. We have integrated the three methods by training TT and ZEPPI so that they predict LRs that can be integrated with PrePPI SM^LR^ in a Bayesian framework. We find that, as measured by ROC and PR curves, PrePPI performs somewhat better than TT (Table1) but it should be recalled that PrePPI was trained on HINT-human and tested on HINT-*E. coli*, while TT was trained on STRING-physical for human. Since STRING defines PPIs between proteins in the same complex as physical, this may degrade its performance on a database primarily containing binary interactions. The integrated Int^LR^ yields the best performance, particularly as measured by its AUPR value.

The main improvement resulting from method integration is a dramatic increase in coverage as can be seen in Figure 2. At FPR ≤ 0.001, the number of predicted PPIs range from about 10,000 for SM^LR^ to close to 20,000 for Int^LR^. Of note, these values are comparable to the estimate of 10,000 PPIs based on an analysis of Y2H data^35^.

In order to gain an additional perspective on our predicted PPIs, we tested whether they produce reliable AF-based models. 400 PPIs were chosen for structure prediction with AF3Complex which is more computationally efficient than AF3 and exhibits somewhat better performance. 312 (78%) of the 400 PPIs yielded models with an interface score ≥ 0.38, the threshold for a reliable model^18^. Of note, SM^LR^ alone (FPR ≤ 0.005) yielded 300 reliable AF3 models while TT^prob^ (with probability > 0.5)^15^ produced 291 reliable models. The two sets share 279 reliable models in common illustrating the synergism of the two approaches. It is not clear at this point whether PrePPI models, on average, are as accurate as AF3 models; most of them are very similar while others overlap substantially in interfacial regions, but the domains adopt altered orientations and, in some cases, the monomer conformations have changed somewhat relative to the PDB template structures used by PrePPI.

In order to gain a proteome-wide perspective of the interactomes introduced in this work, we clustered PPIs into subnetworks. Although no functional information was explicitly included in SM, TT, and Int scoring, the clusters were found to have functional coherence as indicated by the Gene Ontology biological process (GO:BP) terms statistically overrepresented for most clusters^23,24^. Although the data presented here relates to the Int^LR^ interactome, we find that SM^LR^ alone produces comparably meaningful clusters whereas TT^prob^ produces fewer statistically significant clusters. Overall, the results of our clustering study provide additional evidence as to the reliability of our PPI predictions. Moreover, they offer a unique proteome-wide perspective of functional relationships on an unprecedented scale including the annotation of proteins previously designated as of unknown function.

## Methods

All calculations were performed for the 4,402 *E. coli K12* (taxonomy identifier 83333) proteins from the UniProtKB reference proteome with identifier UP000000625^36^.

### PrePPI SM^LR^

The PrePPI algorithm has been described multiple times in the literature^9, 10, 12^ and its main features are summarized above. Although PrePPI incorporates many features that enable the prediction of PPIs that do not involve direct physical interactions, here we focus on the structural modeling component of PrePPI, SM, which predicts the physical interaction of two monomeric structured domains. PrePPI’s rapid SM scoring function allows us to evaluate billions of domain-domain and full length protein-protein interactions. Bayesian statistics are used to calculate a likelihood ratio, SM^LR^, for a given putative PPI. This requires the use of true positive (TP) and true negative (TN) databases. The most recent version of SM^LR^ was trained on the human HINT literature-curated high-quality binary database^20^ as the TP set and any PPI not functionally annotated in any experimentally derived databases as the TN set^8^. Out of the 9.6 million potential protein-protein pairwise combinations in *E. coli*, PrePPI is able to evaluate and make predictions for 5.6 million. Of these, the 38,700 PPIs associated with an FPR ≤ 0.005 appear in the online database for *E. coli*, but all entries are available for download.

### Integrating ZEPPI with PrePPI

ZEPPI^16^ leverages co-evolution and conservation signals across protein-protein interfacial contacts to compute a Z-score based on comparisons to randomly constructed interfaces derived from residue pairs found anywhere on a protein surface. ZEPPI relies on species-matched paired multiple sequence alignments which are compiled using Jackhmmer 2.0^37^ to search for homologous sequences for each protein in the EggNog 5.0 database^38^. In all, we were able to calculate effective ZEPPI scores for 5M PPIs. ZEPPI can be used to evaluate any complex whose atomic coordinates are known but here we incorporate it directly into PrePPI as a separate source of evidence. In order to incorporate ZEPPI into a Bayesian network, we converted ZEPPI Z-values into LRs defined here as ZEPPI^LR^. This is accomplished by training ZEPPI Z-scores on HINT for *E. coli*.

### Integrating Topsy-Turvy with PrePPI

Topsy-Turvy^15^, TT, is a sequence-based method to predict PPIs from the Berger/Cowan labs. TT was developed following the earlier introduction of D-SCRIPT^14^, which is based on a protein language model. TT also leverages global “top-down” methods that infer properties from patterns of known PPIs. We applied TT to all *E. coli* sequences resulting in 9.5 million predicted PPIs. Like PrePPI, TT is extremely fast and can applied proteome-wide. It outputs interaction probabilities which, for integration with PrePPI, must be converted to LRs. TT was trained on the STRING^17^ physical database for human. We converted PPI probabilities into LRs by training on the same data set. We define TT^prob^ as containing interaction probabilities obtained from the published version of TT while TT^LR^ provides LRs from our additional stage of training.

### Bayesian model

TT^LR^ was combined with PrePPI and ZEPPI to yield “Int” which integrates all three evidence sources used in this work.

Int^LR^ = SM^LR^ · ZEPPI^LR^ · TT^LR^

During training, ZEPPI scores and TT probabilities are divided into bins using Doane’s formula for automatic binning^39^. 38 bins are used for ZEPPI and 38 for TT. Doane’s formula adjusts the bin count based on skewness. This makes it more suitable for non-normally distributed data and large sample sizes. Since there are 5.6 million, 5 million and 9.6 million predicted PPIs for SM^LR^, ZEPPI^LR^ and TT^LR^, respectively, there will be many PPIs for which no value can be calculated for SM^LR^ and/or ZEPPI^LR^. In these cases, the corresponding LRs are set to 1 when ROC and PR curves are calculated. Thus, a large fraction of our calculated LRs are based entirely on TT.

### Choice of PPIs to be studied with AF3Complex

To relate our PPI predictions to models constructed with AF3Complex, we created a set of 400 Int^LR^ PPIs with FPRs ≤ 0.001 based on the ROC curves obtained from testing on HINT (Figure 1). There are 20,523 protein-protein interactions above this cutoff involving 3,219 proteins. To select a challenging subset, we retained only those pairs whose members have low local sequence identity (< 40%) to and sufficient alignment length (> 100 residues) with the corresponding chains from the PDB template complex selected by PrePPI for model evaluation. This filtering step decreased the number of PPIs from 20,523 to 7,421. Further, to avoid overrepresentation, no protein appears more than six times in this set. From the highest-confidence PPIs matching these criteria, 400 were chosen so that half appear in databases (and, thus, half are novel predictions) and 60 are homodimeric. Detailed information on the final set of 400 PPIs is provided in Supplementary Table 1A. Note that all 400 PPIs are positive predictions in the sense that the Int^LR^ is always high (FPR ≤ 0.001).

### Interactome clustering and functional annotation

The Markov clustering algorithm as implemented in Cytoscape^19, 22^ was used to cluster the network obtained from the Int^LR^ PPI predictions with FPR ≤ 0.001 (Figure 1A). Default parameters were used and all edge weights set to 1. Over-Representation Analysis for Gene Ontology (GO) terms (biological process (BP), cellular compartment (CC), and molecular function (MF)) was performed with the enrichGO function in the clusterProfiler R package (v4.14.6)^24^. The org.EcK12.eg.db R package^39^ was used for *E. coli strain K12* genome-wide annotation with gene symbols. Enriched GO terms with an adjusted p-value ≤ 0.05 were retained. The adjusted p-value was determined with the Benjamini-Hochberg (BH) method to correct for multiple hypothesis testing. Network, clustering and annotation results are provided in Supplementary Table S2.

## Figure and Table Legends

**Figure S1.**
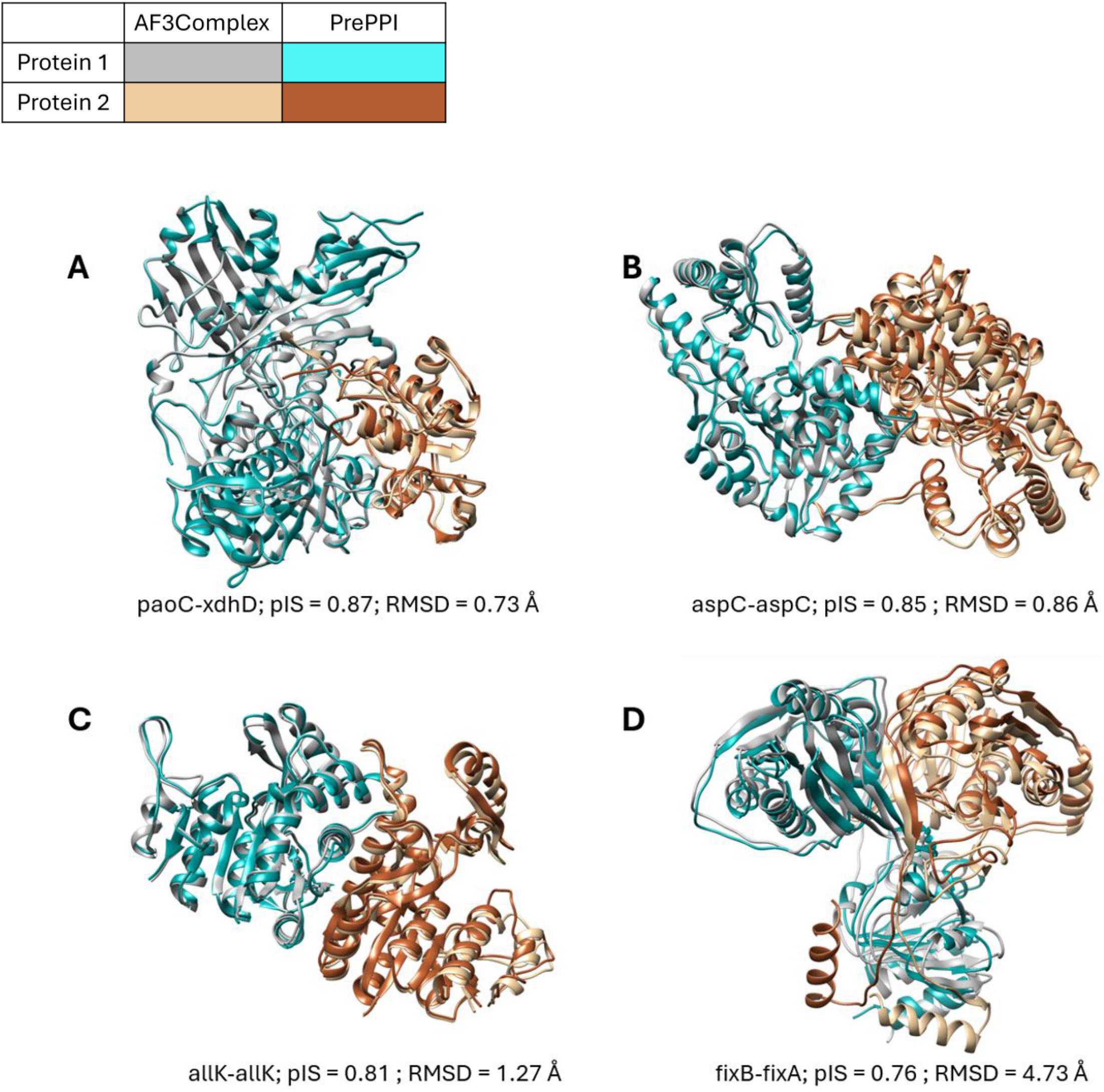
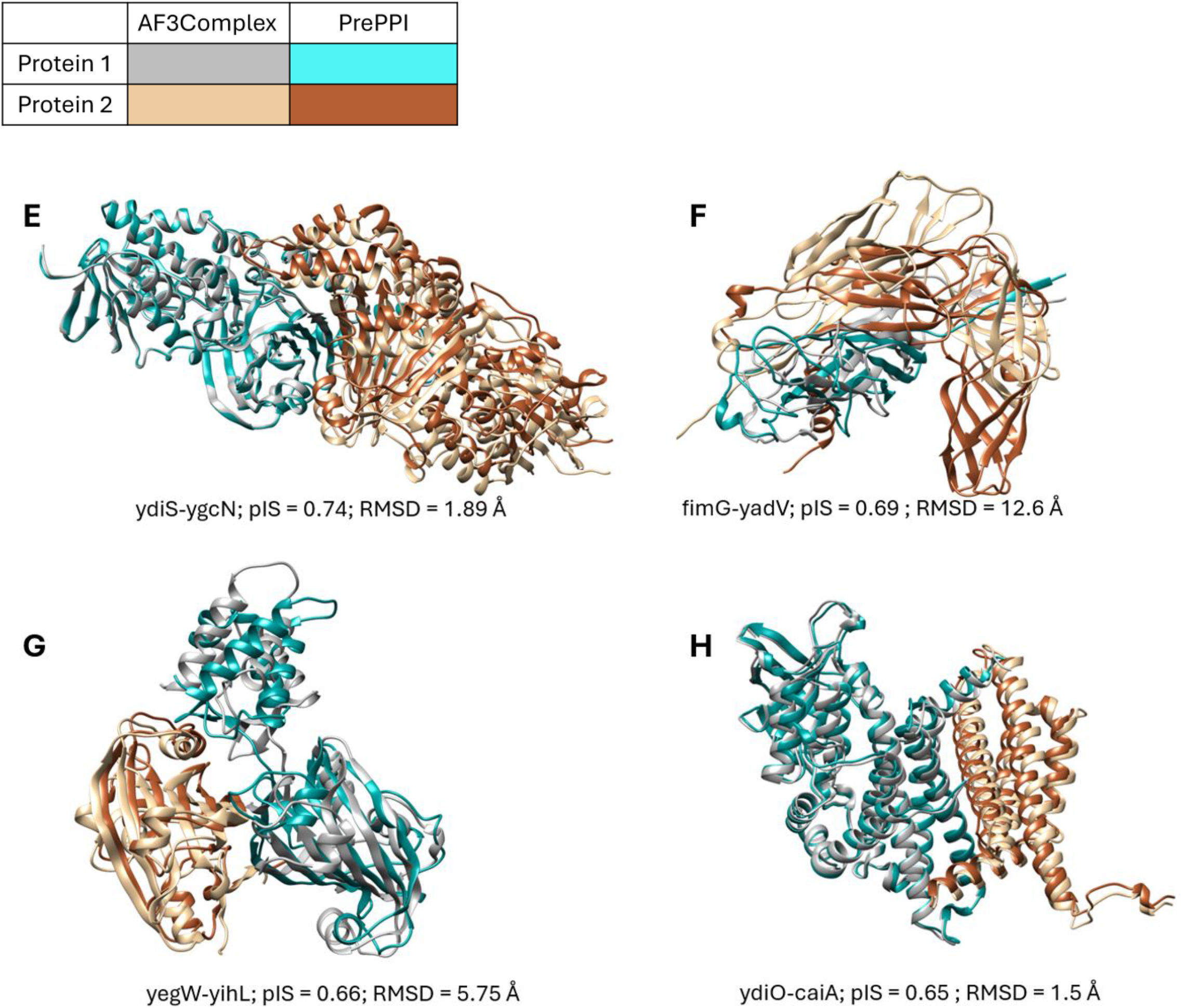
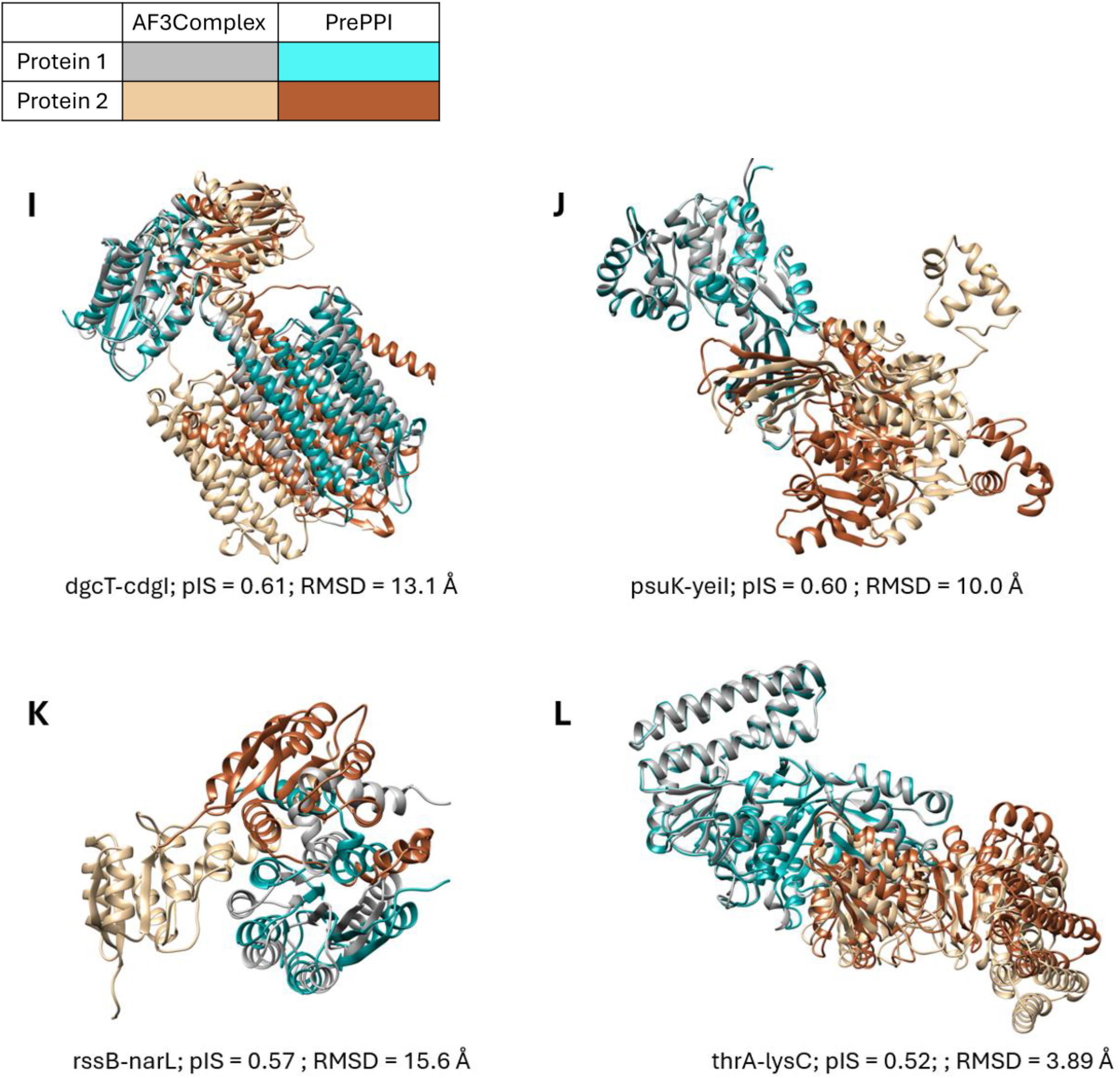
Superposition of PrePPI and AF3Complex models. Models are represented in backbone ribbon representation. The same AF3Complex models from Figure 3 are depicted (gray and gold) superimposed with the PrePPI models for the same PPI (cyan and brown). Proteins are denoted by their gene names; pIS is the predicted interface score for the AF3Complex models; and RMSD is from the superposition of the AF3Complex and PrePPI models. Information on the superpositions is provided in Supplementary Table S1B. **A-D**. Superposition of PrePPI and AF3Complex models. **E-H**. Superposition of PrePPI and AF3Complex models. **I-L**. Superposition of PrePPI and AF3Complex models.

## Acknowledgements

This research was supported in part by grants, R35-GM139585 (BH) and R35-GM118039 (JS) from the Division of General Medical Sciences of the National Institutes of Health.

## Author Contributions

BH, HZ, DM, designed research, analyzed results, and wrote the manuscript. HZ, DM, CV, AN, performed research. AS contributed software tools. JF and JS contributed AF3Complex models and analysis.

## Data availability

High confidence (FPR ≤ 0.005) predictions for PrePPI^SM^ will be available upon publication.

## References

1. Jumper J, Evans R, Pritzel A, Green T, Figurnov M, Ronneberger O, Tunyasuvunakool K, Bates R, Zidek A, Potapenko A, Bridgland A, Meyer C, Kohl SAA, Ballard AJ, Cowie A, Romera-Paredes B, Nikolov S, Jain R, Adler J, Back T, Petersen S, Reiman D, Clancy E, Zielinski M, Steinegger M, Pacholska M, Berghammer T, Bodenstein S, Silver D, Vinyals O, Senior AW, Kavukcuoglu K, Kohli P, Hassabis D. Highly accurate protein structure prediction with AlphaFold. Nature. 2021;596(7873):583–9. Epub 2021/07/16. doi: 10.1038/s41586-021-03819-2. PubMed PMID: 34265844; PMCID: PMC8371605.

2. Abramson J, Adler J, Dunger J, Evans R, Green T, Pritzel A, Ronneberger O, Willmore L, Ballard AJ, Bambrick J, Bodenstein SW, Evans DA, Hung CC, O’Neill M, Reiman D, Tunyasuvunakool K, Wu Z, Zemgulyte A, Arvaniti E, Beattie C, Bertolli O, Bridgland A, Cherepanov A, Congreve M, Cowen-Rivers AI, Cowie A, Figurnov M, Fuchs FB, Gladman H, Jain R, Khan YA, Low CMR, Perlin K, Potapenko A, Savy P, Singh S, Stecula A, Thillaisundaram A, Tong C, Yakneen S, Zhong ED, Zielinski M, Zidek A, Bapst V, Kohli P, Jaderberg M, Hassabis D, Jumper JM. Accurate structure prediction of biomolecular interactions with AlphaFold 3. Nature. 2024;630(8016):493–500. Epub 2024/05/09. doi: 10.1038/s41586-024-07487-w. PubMed PMID: 38718835; PMCID: PMC11168924 including 63/611,674, 63/611,638 and 63/546,444 relating to predicting 3D structures of molecule complexes using embedding neural networks and generative models. All of the authors other than A.B., Y.A.K. and E.D.Z. have commercial interests in the work described.

3. Evans R, O’Neill M, Pritzel A, Antropova N, Senior A, Green T, Žídek A, Bates R, Blackwell S, Yim J, Ronneberger O, Bodenstein S, Zielinski M, Bridgland A, Potapenko A, Cowie A, Tunyasuvunakool K, Jain R, Clancy E, Kohli P, Jumper J, Hassabis D. Protein complex prediction with AlphaFold-Multimer. bioRxiv. 2022:2021.10.04.463034. doi: 10.1101/2021.10.04.463034.

4. Durham J, Zhang J, Humphreys IR, Pei J, Cong Q. Recent advances in predicting and modeling protein-protein interactions. Trends Biochem Sci. 2023;48(6):527–38. Epub 20230414. doi: 10.1016/j.tibs.2023.03.003. PubMed PMID: 37061423.

5. Cong Q, Anishchenko I, Ovchinnikov S, Baker D. Protein interaction networks revealed by proteome coevolution. Science. 2019;365(6449):185–9. Epub 2019/07/13. doi: 10.1126/science.aaw6718. PubMed PMID: 31296772; PMCID: PMC6948103.

6. Humphreys IR, Pei J, Baek M, Krishnakumar A, Anishchenko I, Ovchinnikov S, Zhang J, Ness TJ, Banjade S, Bagde SR, Stancheva VG, Li XH, Liu K, Zheng Z, Barrero DJ, Roy U, Kuper J, Fernandez IS, Szakal B, Branzei D, Rizo J, Kisker C, Greene EC, Biggins S, Keeney S, Miller EA, Fromme JC, Hendrickson TL, Cong Q, Baker D. Computed structures of core eukaryotic protein complexes. Science. 2021;374(6573):eabm4805. Epub 2021/11/12. doi: 10.1126/science.abm4805. PubMed PMID: 34762488; PMCID: PMC7612107.

7. Zhang J, Humphreys IR, Pei J, Kim J, Choi C, Yuan R, Durham J, Liu S, Choi H-J, Baek M, Baker D, Cong Q. Computing the Human Interactome. bioRxiv. 2024:2024.10.01.615885. doi: 10.1101/2024.10.01.615885.

8. Petrey D, Zhao H, Trudeau SJ, Murray D, Honig B. PrePPI: A Structure Informed Proteome-wide Database of Protein-Protein Interactions. J Mol Biol. 2023;435(14):168052. Epub 2023/03/19. doi: 10.1016/j.jmb.2023.168052. PubMed PMID: 36933822; PMCID: PMC10293085.

9. Garzon JI, Deng L, Murray D, Shapira S, Petrey D, Honig B. A computational interactome and functional annotation for the human proteome. Elife. 2016;5. Epub 2016/10/23. doi: 10.7554/eLife.18715. PubMed PMID: 27770567; PMCID: PMC5115866.

10. Zhang QC, Petrey D, Deng L, Qiang L, Shi Y, Thu CA, Bisikirska B, Lefebvre C, Accili D, Hunter T, Maniatis T, Califano A, Honig B. Structure-based prediction of protein-protein interactions on a genome-wide scale. Nature. 2012;490(7421):556–60. Epub 2012/10/02. doi: 10.1038/nature11503. PubMed PMID: 23023127; PMCID: PMC3482288.

11. Burley SK, Berman HM, Kleywegt GJ, Markley JL, Nakamura H, Velankar S. Protein Data Bank (PDB): The Single Global Macromolecular Structure Archive. Methods Mol Biol. 2017;1607:627–41. Epub 2017/06/03. doi: 10.1007/978-1-4939-7000-1_26. PubMed PMID: 28573592; PMCID: PMC5823500.

12. Lasso G, Mayer SV, Winkelmann ER, Chu T, Elliot O, Patino-Galindo JA, Park K, Rabadan R, Honig B, Shapira SD. A Structure-Informed Atlas of Human-Virus Interactions. Cell. 2019;178(6):1526–41 e16. Epub 2019/09/03. doi: 10.1016/j.cell.2019.08.005. PubMed PMID: 31474372; PMCID: PMC6736651.

13. Broyde J, Simpson DR, Murray D, Paull EO, Chu BW, Tagore S, Jones SJ, Griffin AT, Giorgi FM, Lachmann A, Jackson P, Sweet-Cordero EA, Honig B, Califano A. Oncoprotein-specific molecular interaction maps (SigMaps) for cancer network analyses. Nat Biotechnol. 2021;39(2):215–24. Epub 2020/09/16. doi: 10.1038/s41587-020-0652-7. PubMed PMID: 32929263; PMCID: PMC7878435.

14. Sledzieski S, Singh R, Cowen L, Berger B. D-SCRIPT translates genome to phenome with sequence-based, structure-aware, genome-scale predictions of protein-protein interactions. Cell Syst. 2021;12(10):969–82 e6. Epub 2021/09/19. doi: 10.1016/j.cels.2021.08.010. PubMed PMID: 34536380; PMCID: PMC8586911.

15. Singh R, Devkota K, Sledzieski S, Berger B, Cowen L. Topsy-Turvy: integrating a global view into sequence-based PPI prediction. Bioinformatics. 2022;38(Suppl 1):i264-i72. doi: 10.1093/bioinformatics/btac258. PubMed PMID: 35758793; PMCID: PMC9235477.

16. Zhao H, Petrey D, Murray D, Honig B. ZEPPI: Proteome-scale sequence-based evaluation of protein-protein interaction models. Proc Natl Acad Sci U S A. 2024;121(21):e2400260121. Epub 20240514. doi: 10.1073/pnas.2400260121. PubMed PMID: 38743624; PMCID: PMC11127014.

17. Szklarczyk D, Gable AL, Nastou KC, Lyon D, Kirsch R, Pyysalo S, Doncheva NT, Legeay M, Fang T, Bork P, Jensen LJ, von Mering C. The STRING database in 2021: customizable protein-protein networks, and functional characterization of user-uploaded gene/measurement sets. Nucleic Acids Res. 2021;49(D1):D605-D12. Epub 2020/11/26. doi: 10.1093/nar/gkaa1074. PubMed PMID: 33237311; PMCID: PMC7779004.

18. Feldman J, Skolnick J. AF3Complex Yields Improved Structural Predictions of Protein Complexes. bioRxiv. 2025. Epub 20250303. doi: 10.1101/2025.02.27.640585. PubMed PMID: 40093092; PMCID: PMC11908126.

19. Morris JH, Apeltsin L, Newman AM, Baumbach J, Wittkop T, Su G, Bader GD, Ferrin TE. clusterMaker: a multi-algorithm clustering plugin for Cytoscape. BMC Bioinformatics. 2011;12:436. Epub 20111109. doi: 10.1186/1471-2105-12-436. PubMed PMID: 22070249; PMCID: PMC3262844.

20. Das J, Yu H. HINT: High-quality protein interactomes and their applications in understanding human disease. BMC Syst Biol. 2012;6:92. Epub 2012/08/01. doi: 10.1186/1752-0509-6-92. PubMed PMID: 22846459; PMCID: PMC3483187.

21. Leimkuhler S. The biosynthesis of the molybdenum cofactors in Escherichia coli. Environ Microbiol. 2020;22(6):2007–26. Epub 20200406. doi: 10.1111/1462-2920.15003. PubMed PMID: 32239579.

22. Shannon P, Markiel A, Ozier O, Baliga NS, Wang JT, Ramage D, Amin N, Schwikowski B, Ideker T. Cytoscape: a software environment for integrated models of biomolecular interaction networks. Genome Res. 2003;13(11):2498–504. doi: 10.1101/gr.1239303. PubMed PMID: 14597658; PMCID: PMC403769.

23. Gene Ontology C, Aleksander SA, Balhoff J, Carbon S, Cherry JM, Drabkin HJ, Ebert D, Feuermann M, Gaudet P, Harris NL, Hill DP, Lee R, Mi H, Moxon S, Mungall CJ, Muruganugan A, Mushayahama T, Sternberg PW, Thomas PD, Van Auken K, Ramsey J, Siegele DA, Chisholm RL, Fey P, Aspromonte MC, Nugnes MV, Quaglia F, Tosatto S, Giglio M, Nadendla S, Antonazzo G, Attrill H, Dos Santos G, Marygold S, Strelets V, Tabone CJ, Thurmond J, Zhou P, Ahmed SH, Asanitthong P, Luna Buitrago D, Erdol MN, Gage MC, Ali Kadhum M, Li KYC, Long M, Michalak A, Pesala A, Pritazahra A, Saverimuttu SCC, Su R, Thurlow KE, Lovering RC, Logie C, Oliferenko S, Blake J, Christie K, Corbani L, Dolan ME, Drabkin HJ, Hill DP, Ni L, Sitnikov D, Smith C, Cuzick A, Seager J, Cooper L, Elser J, Jaiswal P, Gupta P, Jaiswal P, Naithani S, Lera-Ramirez M, Rutherford K, Wood V, De Pons JL, Dwinell MR, Hayman GT, Kaldunski ML, Kwitek AE, Laulederkind SJF, Tutaj MA, Vedi M, Wang SJ, D’Eustachio P, Aimo L, Axelsen K, Bridge A, Hyka-Nouspikel N, Morgat A, Aleksander SA, Cherry JM, Engel SR, Karra K, Miyasato SR, Nash RS, Skrzypek MS, Weng S, Wong ED, Bakker E, Berardini TZ, Reiser L, Auchincloss A, Axelsen K, Argoud-Puy G, Blatter MC, Boutet E, Breuza L, Bridge A, Casals-Casas C, Coudert E, Estreicher A, Livia Famiglietti M, Feuermann M, Gos A, Gruaz-Gumowski N, Hulo C, Hyka-Nouspikel N, Jungo F, Le Mercier P, Lieberherr D, Masson P, Morgat A, Pedruzzi I, Pourcel L, Poux S, Rivoire C, Sundaram S, Bateman A, Bowler-Barnett E, Bye AJH, Denny P, Ignatchenko A, Ishtiaq R, Lock A, Lussi Y, Magrane M, Martin MJ, Orchard S, Raposo P, Speretta E, Tyagi N, Warner K, Zaru R, Diehl AD, Lee R, Chan J, Diamantakis S, Raciti D, Zarowiecki M, Fisher M, James-Zorn C, Ponferrada V, Zorn A, Ramachandran S, Ruzicka L, Westerfield M. The Gene Ontology knowledgebase in 2023. Genetics. 2023;224(1). Epub 2023/03/04. doi: 10.1093/genetics/iyad031. PubMed PMID: 36866529; PMCID: PMC10158837.

24. Yu G. Thirteen years of clusterProfiler. Innovation (Camb). 2024;5(6):100722. Epub 20241021. doi: 10.1016/j.xinn.2024.100722. PubMed PMID: 39529960; PMCID: PMC11551487.

25. UniProt C. UniProt: the Universal Protein Knowledgebase in 2023. Nucleic Acids Res. 2022. Epub 2022/11/22. doi: 10.1093/nar/gkac1052. PubMed PMID: 36408920.

26. Garcia-Heredia A. Plasma Membrane-Cell Wall Feedback in Bacteria. J Bacteriol. 2023;205(3):e0043322. Epub 20230216. doi: 10.1128/jb.00433-22. PubMed PMID: 36794934; PMCID: PMC10029715.

27. Munhoz DD, Richards AC, Santos FF, Mulvey MA, Piazza RMF. E. coli Common pili promote the fitness and virulence of a hybrid aEPEC/ExPEC strain within diverse host environments. Gut Microbes. 2023;15(1):2190308. doi: 10.1080/19490976.2023.2190308. PubMed PMID: 36949030; PMCID: PMC10038029.

28. Wu H, Fives-Taylor PM. Molecular strategies for fimbrial expression and assembly. Crit Rev Oral Biol Med. 2001;12(2):101–15. doi: 10.1177/10454411010120020101. PubMed PMID: 11345521.

29. Guo K, Gao H. Physiological Roles of Nitrite and Nitric Oxide in Bacteria: Similar Consequences from Distinct Cell Targets, Protection, and Sensing Systems. Adv Biol (Weinh). 2021;5(9):e2100773. Epub 20210726. doi: 10.1002/adbi.202100773. PubMed PMID: 34310085.

30. Gagarinova A, Hosseinnia A, Rahmatbakhsh M, Istace Z, Phanse S, Moutaoufik MT, Zilocchi M, Zhang Q, Aoki H, Jessulat M, Kim S, Aly KA, Babu M. Auxotrophic and prototrophic conditional genetic networks reveal the rewiring of transcription factors in Escherichia coli. Nat Commun. 2022;13(1):4085. Epub 20220714. doi: 10.1038/s41467-022-31819-x. PubMed PMID: 35835781; PMCID: PMC9283627.

31. Anjou C, Lotoux A, Morvan C, Martin-Verstraete I. From ubiquity to specificity: The diverse functions of bacterial thioredoxin systems. Environ Microbiol. 2024;26(6):e16668. doi: 10.1111/1462-2920.16668. PubMed PMID: 38899743.

32. Huttlin EL, Bruckner RJ, Navarrete-Perea J, Cannon JR, Baltier K, Gebreab F, Gygi MP, Thornock A, Zarraga G, Tam S, Szpyt J, Gassaway BM, Panov A, Parzen H, Fu S, Golbazi A, Maenpaa E, Stricker K, Guha Thakurta S, Zhang T, Rad R, Pan J, Nusinow DP, Paulo JA, Schweppe DK, Vaites LP, Harper JW, Gygi SP. Dual proteome-scale networks reveal cell-specific remodeling of the human interactome. Cell. 2021;184(11):3022–40 e28. Epub 2021/05/08. doi: 10.1016/j.cell.2021.04.011. PubMed PMID: 33961781; PMCID: PMC8165030.

33. Huttlin EL, Ting L, Bruckner RJ, Gebreab F, Gygi MP, Szpyt J, Tam S, Zarraga G, Colby G, Baltier K, Dong R, Guarani V, Vaites LP, Ordureau A, Rad R, Erickson BK, Wuhr M, Chick J, Zhai B, Kolippakkam D, Mintseris J, Obar RA, Harris T, Artavanis-Tsakonas S, Sowa ME, De Camilli P, Paulo JA, Harper JW, Gygi SP. The BioPlex Network: A Systematic Exploration of the Human Interactome. Cell. 2015;162(2):425–40. Epub 2015/07/18. doi: 10.1016/j.cell.2015.06.043. PubMed PMID: 26186194; PMCID: PMC4617211.

34. Luck K, Kim DK, Lambourne L, Spirohn K, Begg BE, Bian W, Brignall R, Cafarelli T, Campos-Laborie FJ, Charloteaux B, Choi D, Cote AG, Daley M, Deimling S, Desbuleux A, Dricot A, Gebbia M, Hardy MF, Kishore N, Knapp JJ, Kovacs IA, Lemmens I, Mee MW, Mellor JC, Pollis C, Pons C, Richardson AD, Schlabach S, Teeking B, Yadav A, Babor M, Balcha D, Basha O, Bowman-Colin C, Chin SF, Choi SG, Colabella C, Coppin G, D’Amata C, De Ridder D, De Rouck S, Duran-Frigola M, Ennajdaoui H, Goebels F, Goehring L, Gopal A, Haddad G, Hatchi E, Helmy M, Jacob Y, Kassa Y, Landini S, Li R, van Lieshout N, MacWilliams A, Markey D, Paulson JN, Rangarajan S, Rasla J, Rayhan A, Rolland T, San-Miguel A, Shen Y, Sheykhkarimli D, Sheynkman GM, Simonovsky E, Tasan M, Tejeda A, Tropepe V, Twizere JC, Wang Y, Weatheritt RJ, Weile J, Xia Y, Yang X, Yeger-Lotem E, Zhong Q, Aloy P, Bader GD, De Las Rivas J, Gaudet S, Hao T, Rak J, Tavernier J, Hill DE, Vidal M, Roth FP, Calderwood MA. A reference map of the human binary protein interactome. Nature. 2020;580(7803):402–8. Epub 2020/04/17. doi: 10.1038/s41586-020-2188-x. PubMed PMID: 32296183; PMCID: PMC7169983.

35. Rajagopala SV, Sikorski P, Kumar A, Mosca R, Vlasblom J, Arnold R, Franca-Koh J, Pakala SB, Phanse S, Ceol A, Hauser R, Siszler G, Wuchty S, Emili A, Babu M, Aloy P, Pieper R, Uetz P. The binary protein-protein interaction landscape of Escherichia coli. Nat Biotechnol. 2014;32(3):285–90. Epub 2014/02/25. doi: 10.1038/nbt.2831. PubMed PMID: 24561554; PMCID: PMC4123855.

36. Breuza L, Poux S, Estreicher A, Famiglietti ML, Magrane M, Tognolli M, Bridge A, Baratin D, Redaschi N, UniProt C. The UniProtKB guide to the human proteome. Database (Oxford). 2016;2016. Epub 20160220. doi: 10.1093/database/bav120. PubMed PMID: 26896845; PMCID: PMC4761109.

37. Eddy SR. Accelerated Profile HMM Searches. PLoS Comput Biol. 2011;7(10):e1002195. Epub 20111020. doi: 10.1371/journal.pcbi.1002195. PubMed PMID: 22039361; PMCID: PMC3197634.

38. Huerta-Cepas J, Szklarczyk D, Heller D, Hernandez-Plaza A, Forslund SK, Cook H, Mende DR, Letunic I, Rattei T, Jensen LJ, von Mering C, Bork P. eggNOG 5.0: a hierarchical, functionally and phylogenetically annotated orthology resource based on 5090 organisms and 2502 viruses. Nucleic Acids Res. 2019;47(D1):D309-D14. Epub 2018/11/13. doi: 10.1093/nar/gky1085. PubMed PMID: 30418610; PMCID: PMC6324079.

39. Carlson M. Genome wide annotation for E coli strain K12. doi: 10.18129/B9.bioc.org.EcK12.eg.db.

40. Doane DP. Aesthetic frequency classifications. American Statistician. 1976;30(4):181–183. doi: 10.1080/00031305.1976.10479172.

